# FABP5 regulates ether lipid metabolism to ameliorate atopic dermatitis

**DOI:** 10.1101/2025.10.24.684205

**Authors:** Mathias H. Skadow, Walter K. Mowel, Holly N. Blackburn, Matthew Z. Madden, Haris Mirza, Fengrui Zhang, Autumn G. York, Richard A. Flavell

## Abstract

Atopic dermatitis is an allergic skin disease associated with a profound reorganization of the epidermal lipidome. The effect of the altered lipidome on the skin-resident immune cells that drive disease is unclear. Previous reports identified Fatty acid binding protein 5 (FABP5) as a biomarker for atopic dermatitis, yet how FABP5 might contribute to disease pathogenesis is unknown. Here, we use a murine model of atopic dermatitis, to demonstrate that FABP5 is highly expressed in immune and epithelial cell lineages and that FABP5 protects against skin inflammation. Lipidomic analysis revealed that FABP5 deficiency broadly disrupts the systemic abundance of ether-linked lipids, a minor but important subset of glycerophospholipids. We show that these changes in ether lipid abundance are crucial for the proper regulation of platelet activating factor (PAF), a potent inflammatory ether lipid derivative. Concordantly, we observe elevated PAF in FABP5-deficient mice with dermatitis and that depletion of basophils, a major source of PAF, is sufficient to ameliorate disease in these animals. Altogether, our findings reveal a novel role for FABP5 in the control of allergic inflammation through the modulation of ether lipid and PAF metabolism.

## Introduction

Atopic dermatitis (AD) is an increasingly common and often debilitating allergic inflammation of the skin, affecting an estimated 10.8% of children in the United States^1,2^. The emergence of atopic dermatitis often precedes the diagnosis of other allergic diseases – asthma, allergic rhinitis, and food allergy – in sequential diagnoses known as atopic march^3,4^. This clinical trajectory underscores the importance of understanding the immunological and tissue-specific mechanisms that drive AD and contribute to the broader development of allergic inflammation.

One important characteristic of AD is the dysregulation of lipid metabolism in the skin^5–8^. These alterations in lipid metabolism can be detected even before the onset of symptoms and can serve as predictive biomarkers of future disease^9,10^. Changes in the composition of the skin lipidome are hypothesized to affect barrier function and may be both a cause and effect of allergic inflammation in the skin^5^. It is now clear that the specific metabolic milieu of different tissues has profound effects on immune cell function^11^. The specific lipids important for forming the skin barrier – ceramides, long-chain fatty acids, and cholesterol – have also been implicated in tailoring the activation of tissue-resident immune cells in other contexts^12–14^. However, the extent to which keratinocytes and immune cells alter the skin lipidome and the functional consequences of these changes remains unclear.

FABPs are a family of cytosolic lipid binding proteins implicated in the control of lipid uptake, fatty acid oxidation, and transcriptional control of lipid metabolic programs through facilitation of nuclear hormone receptor signaling^15^. These proteins therefore represent a central node in the coordination of lipid metabolism. While FABPs are abundant cytosolic proteins, they are also observed in circulation, and their role in the extracellular space is currently being explored^16–18^. One paralog, FABP5, is highly expressed in keratinocytes, but also displays wide tissue distribution, including expression in lymphoid tissues, lung, and central nervous system^15^. FABP5 has been previously demonstrated to ameliorate allergic inflammation in several mouse models, though reports differ regarding its mechanistic role^19,20^. Interestingly, AD has been associated with increased concentrations of FABP5 in the skin and serum of patients^21^, but how FABP5 may affect lipid metabolism during disease and whether it contributes to disease severity remains unclear.

In this study we utilized a mouse model of AD to demonstrate that FABP5 is upregulated in skin and circulation during disease to constrain inflammation. Moreover, we found that FABP5 is expressed by multiple immune and non-immune cell types during AD, and expression in either lineage is sufficient to constrain inflammation. Unexpectedly, we found that FABP5 is crucial for the proper metabolism of ether-linked lipids and that FABP5 deficiency leads to an increase in platelet activating factor (PAF) an important ether lipid derivative. Finally, we demonstrated that we could ameliorate disease severity in these animals through the therapeutic alteration of PAF metabolism or depletion of basophils, an abundant PAF-secreting cell type, highlighting the importance of this pathway in AD.

### FABP5 expression is upregulated during dermatitis and constrains disease

AD and atopic march have been previously associated with increased FABP5 expression in patient lesional skin and serum^21^. To interrogate whether similar patterns of FABP5 expression could be observed in murine AD, we re-analyzed bulk RNAseq data^22^ comparing mice treated with vehicle or MC903, a synthetic vitamin D analog that drives expression of *Tslp* in keratinocytes to induce AD-like disease^23^. Consistent with previous observations in AD patients, we found that MC903 treatment resulted in increased *Fabp5* expression in mouse skin but did not induce the expression of other FABP family members, including the closely related gene, *Fabp4* (Figure 1A). Congruent with this finding, we observed dramatically increased FABP5 transcript and protein levels in the skin of mice with MC903-induced dermatitis compared to vehicle treated controls (Figure 1B, 1C). Importantly, we also observed that MC903 treatment resulted in a marked increase in the concentration of circulating FABP5 (Figure 1D). Together, these data demonstrate that FABP5 accumulates in the skin and circulation downstream of MC903-induced AD, implying that FABP5 may play a role in allergic inflammation.

**Figure 1:**
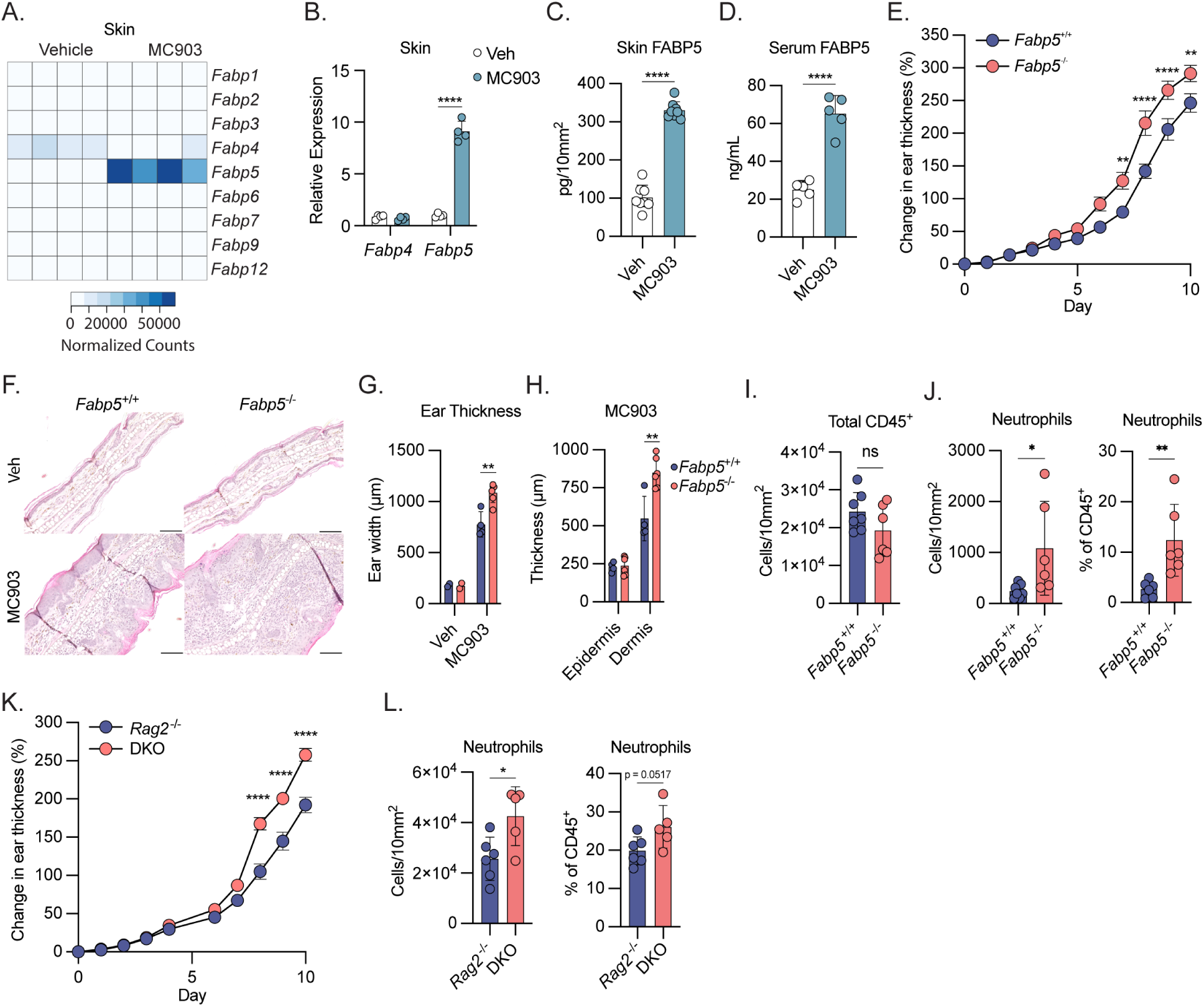
FABP5 is upregulated during dermatitis and ameliorates disease. A) Heat map of gene expression from RNAseq of naïve (vehicle treated) and dermatotic (MC903 treated) mouse skin showing expression of all FABP family members. Color is scaled by normalized transcript counts. B) qPCR analysis of *Fabp4* and *Fabp5* gene expression from mouse vehicle or MC903-treated mouse skin. (n = 8) C) Quantification of FABP5 protein via ELISA from mouse skin punch biopsies collected on day 10 of treatment with vehicle or MC903. Protein is normalized per unit area of skin. (n = 16) D) Quantification of FABP5 protein via ELISA from mouse serum from day 10 of treatment with vehicle or MC903-treated mice. (n = 10) E) Caliper measurements showing increase in ear thickness over time during MC903 treatment of *Fabp5^+/+^*and *Fabp5^-/-^* mice (n = 14). Dots show mean ± SEM. Stars represent Šidák corrected *p* values for 2-way ANOVA with multiple comparisons. F) Representative images from H&E histology showing transverse sections of ear pinnae from vehicle and MC903-treated *Fabp5^+/+^* and *Fabp5^-/-^* mice. G) Quantification of ear thickness based on histological measurements of transverse section of ear pinnae. (vehicle n = 4; MC903 n = 10) H) Quantification of epidermal and dermal thickness based on histological measurements of transverse section of ear pinnae. (vehicle n = 4; MC903 n = 10) I) Flow cytometry-based quantification of total CD45^+^ cells in the skin of MC903-treated *Fabp5^+/+^* and *Fabp5^-/-^* mice. Cell count is normalized per unit area. (n = 13) J) Flow cytometry-based quantification of total neutrophils in the skin of MC903-treated *Fabp5^+/+^*and *Fabp5^-/-^* mice. Cell count is normalized per unit area (left) or reported as a percent of total CD45^+^ immune cells (right). (n = 13) K) Caliper measurements showing increase in ear thickness over time during MC903 treatment of Rag2^-/-^ and *Fabp5^-/-^ Rag2^-/-^* (DKO) mice (n = 14). Dots show mean ± SEM. Stars represent Šidák corrected *p* values for 2-way ANOVA with multiple comparisons. L) Flow cytometry-based quantification of total neutrophils in the skin of *Rag2^-/-^* and *Fabp5^-/-^ Rag2^-/-^* (DKO) mice on day 10 of MC903 treatment. Cell count is normalized per unit area (left) or reported as a percent of total CD45^+^ immune cells (right). (n = 11) Unless otherwise noted, all data are reported as means ± SD. * *p* < 0.05; ** *p* < 0.01, *** *p* < 0.005. Two-tailed unpaired Student’s *t*-test. Datapoints are discrete biological replicates.

Previous work has proposed that FABP5 may exacerbate dermatitis^21^, however other literature demonstrates a role for FABP5 in constraining allergic inflammation^19,20^. To definitively test the role of FABP5 during dermatitis, we created *Fabp5^-/-^*mice by excising the *Fabp5* coding region on mouse chromosome 3. FABP5 protein could not be detected in serum of Fabp5 KO mice validating the knockout strategy (data not shown). To test the role of FABP5 in AD, we challenged FABP5 KO mice and their control littermates with MC903 and measured skin thickness to track disease progression. Consistent with previous results, daily MC903 challenge induced robust skin inflammation and skin thickening in wild-type controls (Figure 1E, 1F). Strikingly, *Fabp5*^-/-^ mice displayed more severe dermatitis apparent in the increase in skin thickness over time and under histological examination (Figure 1E, 1F). Importantly, we observed no obvious effects of FABP5 deficiency on skin morphology at steady state (Figure 1F). Further histological analysis revealed that while both genotypes displayed a similar magnitude of epidermal hyperplasia following MC903 treatment (Figure 1H), FABP5-deficient mice had a significant increase in dermal thickness, which explained the observed overall increase in skin thickness (Figure 1H).

In AD, infiltrating innate immune cells, such as neutrophils, monocytes, eosinophils and basophils, are recruited to the skin by activated endothelial cells and local production of chemokines^24^. To test whether the increase in dermal thickness in FABP5-deficient mice may be coincident with increased immune cell infiltration, we isolated immune cells from the skin of both WT and FABP5 KO animals treated with MC903 for profiling via flow cytometry (See Figure S1A-C for gating strategy). Surprisingly, despite the dramatic increase in dermal thickness, immune profiling of the skin did not reveal an increase in total infiltrating CD45^+^ immune cells (Figure 1I). Indeed, we found that most innate immune cell populations that infiltrate the dermis were not differentially abundant between WT and FABP5-KO animals (Figure S2A). Instead, we observed a specific increase in the number and frequency of skin-infiltrating neutrophils (Figure 1J).

FABP5 (in coordination with FABP4), has been shown to be important for lipid uptake and survival of T cells in tissues^25^. Moreover, FABP5 has been demonstrated to be a key regulator of regulatory T cell suppressor function, and through its ability to modulate PPARγ signaling may affect Th17 differentiation^26,21,27^. The pathology of MC903-induced dermatitis does not rely on the adaptive immune system^28^. Therefore, to rule out any potential effects of FABP5 in the T cell compartment, we crossed *Fabp5*^-/-^ mice to a *Rag*2^-/-^ background which are unable to produce T or B cells. We found that *Fabp5*^-/-^*Rag*2^-/-^ double knockout animals developed worse dermatitis than their *Fabp5*^+/+^*Rag*2^-/-^ littermate controls (Figure 1K) FABP5-deficiency in Rag2^-/-^ mice similarly resulted in an increase in skin-infiltrating neutrophils (Figure 1L). Consistent with these findings, we found that CD4^+^ and CD8^+^ T cell numbers were either comparable or slightly increased in the skin of FABP5 KO mice compared to WT controls after MC903 treatment (Figure S2B). We also examined the polarization of T helper cells in the skin after MC903 treatment and found that FABP5 deficiency did not affect the abundance of different T helper subsets (Figure S2C). Together, these data suggest that FABP5 acts independently of the adaptive immune system to constrain dermatitis severity and limit neutrophil infiltration in the skin during AD.

### FABP5 is expressed and secreted by multiple cell types during MC903-induced dermatitis

FABP5, previously known as epidermal-FABP (E-FABP), is expressed highly in keratinocytes. However, FABP5 is also highly expressed in various immune cell populations, endothelial cells, and adipocytes, all of which could contribute to the pathology of atopic dermatitis. To test which cell types upregulate *Fabp5* during dermatitis, we performed single cell RNA sequencing (scRNAseq) on cells isolated from skin of RAG2-KO mice treated with MC903 or vehicle control. UMAP analysis resolved multiple clusters, including keratinocytes, innate lymphoid cells (ILCs), basophils, mast cells, and multiple monocyte, macrophage and dendritic cell (DC) populations (Figure 2A, 2B). *Fabp5* expression was markedly upregulated in keratinocytes, endothelial cells, and multiple myeloid populations following MC903 treatment (Figure 2C), suggesting that upregulation of Fabp5 may be a shared response downstream of type 2 inflammation.

**Figure 2:**
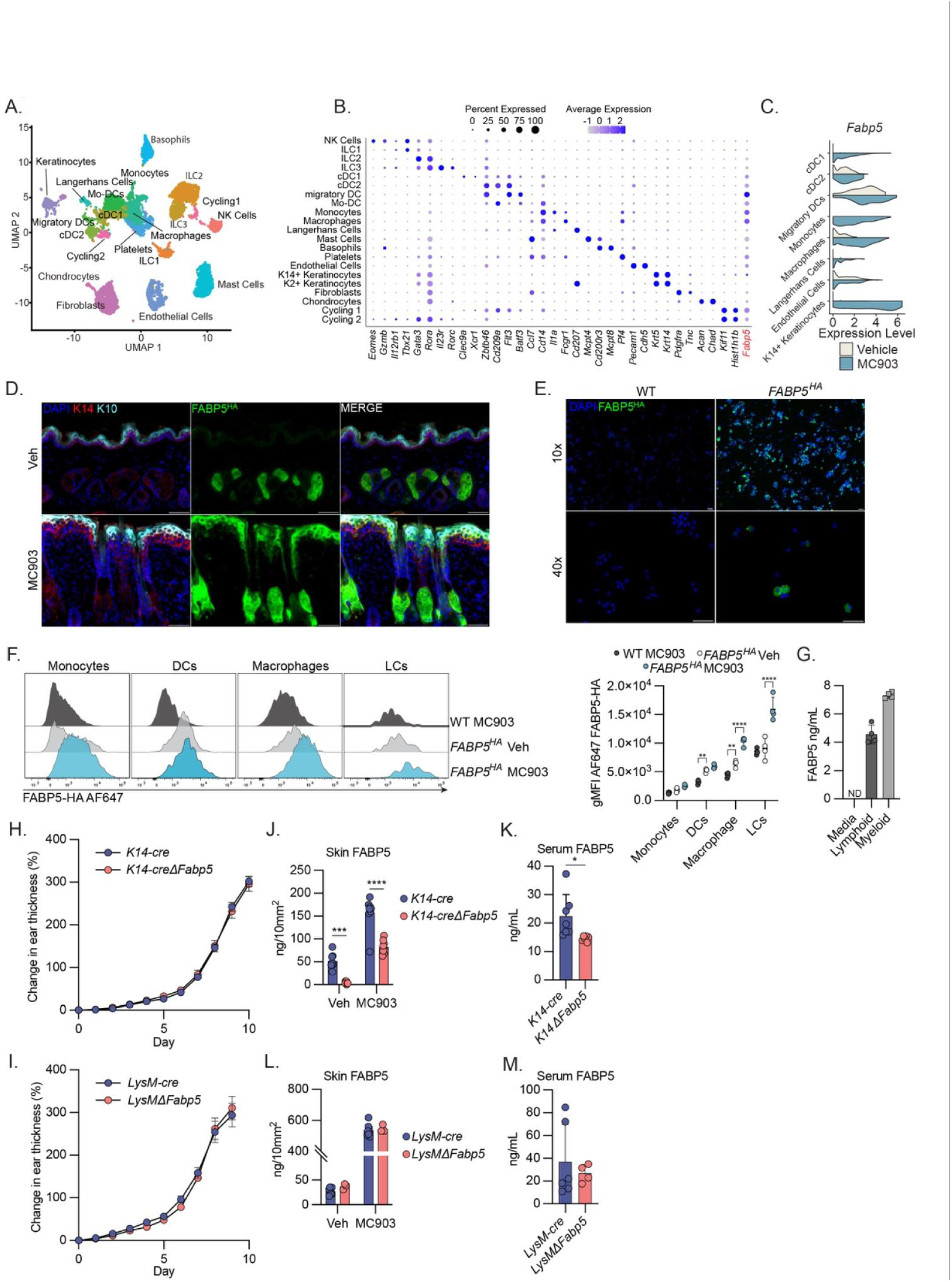
FABP5 is expressed by multiple cell types to constrain disease. A) UMAP analysis showing manually annotated cell clusters from skin of *Rag2^-/-^* mice treated with vehicle or MC903. B) Dot plot visualization of expression of select genes used in identifying cell clusters in (A). C) Split violin plots displaying gene expression of Fabp5 in select cell clusters from (A), comparing vehicle treated (white) and MC903 treated (blue). D) Representative immunofluorescence images of skin from FABP5^HA^ mice on day 10 of treatment with vehicle or MC903. E) Low and high magnification immunofluorescence images of CD45^+^ immune cells isolated from skin of wild-type (WT) or FABP5^HA^ mice on day 10 of treatment with MC903. F) Histograms (left) comparing the fluorescent intensity of intracellular α-ΗΑ AF647 staining in monocytes, dendritic cells (DCs), macrophages, and Langerhans cells (LCs) isolated from skin of WT and FABP5^HA^ mice treated with vehicle or MC903. Quantification of geometric mean fluorescent intensity (right) from these same populations (n = 12). G) Quantification via ELISA of FABP5 in cell culture supernatants from CD45+CD90+ (Lymphoid) or CD45+CD11b+Ly6G-(Myeloid) cells isolated from MC903-treated skin or media only control (n = 8 technical replicates) H) Caliper measurements showing increase in ear thickness over time during MC903 treatment of *K14-cre* and *K14-creFabp5^Fl/Fl^* (*K14ΔFabp5*) mice (n = 18). Dots show mean ± SEM. I) Caliper measurements showing increase in ear thickness over time during MC903 treatment of *LysM-cre* and *LysM-creFabp5^Fl/Fl^* (*LysMΔFabp5*) mice (n = 10). Dots show mean ± SEM. J) Quantification via ELISA of FABP5 from skin punch biopsies taken from of *K14-cre* and *K14-creFabp5^Fl/Fl^*(*K14ΔFabp5*) mice on day 10 of treatment with vehicle or MC903. Protein abundance is normalized to area of skin. (n = 16) K) Quantification via ELISA of FABP5 protein in serum of *K14-cre* and *K14-creFabp5^Fl/Fl^* (*K14ΔFabp5*) mice on day 10 of treatment with MC903. (n = 13) L) Quantification via ELISA of FABP5 protein from skin punch biopsies taken from of *LysM-cre* and *LysM-creFabp5^Fl/Fl^* (*LysMΔFabp5*) mice on day 10 of treatment with vehicle or MC903. Protein abundance is normalized by punch biopsy area. (n = 11) M) Quantification via ELISA of FABP5 protein in serum of of *LysM-cre* and *LysM-creFabp5^Fl/Fl^* (*LysMΔFabp5*) mice on day 10 of treatment with MC903. (n = 10) Unless otherwise noted, all data are reported as means ± SD. * *p* < 0.05; ** *p* < 0.01, *** *p* < 0.005. Two-tailed unpaired Student’s *t*-test. Datapoints are discrete biological replicates.

To further confirm these findings and to obtain spatial resolution of FABP5 expression during disease, we generated FABP5^HA^ mice in which a sequence encoding the HA epitope tag was inserted into the N-terminus of the endogenous *Fabp5* locus. In congruence with previous findings^29^, immunofluorescence microscopy of the skin of naïve mice revealed that FAPB5^HA^ expression in the skin is largely restricted to sebaceous glands during homeostasis (Figure 2D). During MC903-induced dermatitis, we observed that FABP5^HA^ expression was still observed in the sebaceous glands, which become enlarged during disease, but was also observed more broadly throughout the epidermis and colocalized with fully differentiated, keratin-10 expressing keratinocytes (Figure 2D). In addition to this robust induction of FABP5 in keratinocytes, we observed high, but heterogeneous expression of FABP5^HA^ in CD45^+^ immune cells isolated from the skin following MC903 challenge (Figure 2E). We confirmed FABP5 expression in immune cells by flow cytometric analysis and observed significant FABP5-HA in skin-resident macrophages and DCs during homeostasis (Figure 2F, S3A, S3B). In response to MC903 treatment, skin-resident macrophages, and Langerhans cells, displayed a significant increase in intracellular FABP5 (Figure 2F). Despite previously published roles for FABP5 in T cells, we observed relatively little FABP5 expression in skin infiltrating CD4+ T cells (Figure S3C). Together, these data demonstrate that multiple myeloid and non-immune cell types induce FABP5 expression in response to MC903-induced dermatitis.

FABP5 is found in circulation in naïve mice and the circulating concentration of FABP5 increases during dermatitis (Figure 1D). Much of the literature presumes that FABPs enter circulation due to nonspecific, cytosolic leak associated with tissue damage rather than from regulated secretion^15^. For this reason, circulating FABP5 is often considered as a biomarker of tissue damage, and potential roles as a cytokine or extracellular regulator of lipid metabolism are overlooked. We sought to determine whether skin-infiltrating immune cell populations were capable of secreting FABP5. We isolated lymphoid (CD45+CD90+) and non-neutrophil myeloid (CD45+CD11b+Ly6G-) cells from the skin of MC903-treated mice and cultured them in complete media for 4h. We analyzed the concentration of FABP5 in the supernatant and found that both populations were capable of secreting FABP5 (Figure 2G).

### FABP5 is expressed by multiple cell types to constrain disease

Data from scRNAseq and FABP5^HA^ immunofluorescence revealed that multiple cell types upregulate FABP5 expression during dermatitis. We therefore sought to use conditional genetics to analyze how cells of different lineages contributed to local and systemic circulating levels of FABP5 and to determine which cell types required intrinsic FABP5 expression to constrain disease. We generated *Fabp5* floxed mice by inserting loxP sites flanking the 2^nd^ and 3^rd^ exon of the *Fabp5* locus. We crossed *Fabp5^fl/fl^* mice to *K14-cre* and *LysM-cre* mice to generate mice with a conditional deletion of FABP5 in the epidermis and myeloid lineage respectively (*K14ΔFabp5*, *LysMΔFabp5*). Surprisingly, when treated with MC903, we observed that conditional deletion of FABP5 from neither keratinocytes nor myeloid cells was sufficient to recapitulate the exacerbated dermatitis observed in germline FABP5 KO mice (Figure 2H, 2I).

We further sought to examine the extent to which these different cell lineages (keratinocytes, and myeloid cells) contribute to FABP5 protein concentrations in skin and circulation. As expected, we found that naïve *K14ΔFabp5* mice had nearly undetectable levels of FABP5 in the skin (Figure 2J). However, following MC903-induced dermatitis there was a significant increase in skin FABP5 concentrations in *K14ΔFabp5* mice, though this was still significantly lower than FABP5 concentrations in the MC903-treated *K14-cre* control mice (Figure 2J). These data suggest that keratinocytes are the major source of FABP5 in the naïve skin, but that upon inflammation, other cell types upregulate FABP5 or are recruited to the skin and contribute a significant proportion of the total pool of tissue FABP5. In examining the serum of these mice, we observed that MC903 treated *K14ΔFabp5* mice had significantly less circulating FABP5 compared to *K14-cre* controls (Figure 2K). While significant, this decrease was partial – an approximate 30% decrease in serum concentration of FABP5 in *K14ΔFabp5* mice – suggesting that the epidermis is a significant, but not sole source of circulating FABP5 protein during AD. In contrast, we observed no change in FABP5 concentrations in skin or circulation in *LysMΔFabp5* animals, suggesting that while myeloid cells express and secrete FABP5, they surprisingly do not contribute substantially to the amount of protein in circulation or tissue (Figure 2L, 2M). Altogether, these results indicate that FABP5 is expressed and secreted by multiple cell types in vivo and suggest that FABP5 may ameliorate dermatitis through the regulation of tissue or systemic lipid metabolism rather than through its canonical role of controlling intracellular lipid metabolism.

### FABP5 alters PAF metabolism during dermatitis

The observation that FABP5-deficiency in individual cell lineages is incapable of recapitulating the exacerbated dermatitis observed in the global FABP5-deficient mice, led us to hypothesize that FABP5 may act through multiple cell types to alter systemic lipid metabolism, by for instance, regulating the availability of free fatty acids. To test this hypothesis, we performed shotgun lipidomic analysis on plasma collected from WT and FABP5 KO animals on day 10 of MC903 treatment. Surprisingly, we did not observe alterations in the abundance of total circulating free fatty acids between WT and FABP5 KO animals (Figure S4A), and abundance of most individual free fatty acid species was comparable (Figure S4B). In fact, the plasma lipidomes of WT and FABP5 KO animals were largely identical (Figure S4C) with the notable exception of ether lipids, which were significantly depleted in FABP5 KO animals (Figure 3A). Specifically, FABP5-deficient animals had lower concentrations of total circulating phosphatidylethanolamine (PE) species containing ether-linked 18:0 tails, which was attributable to a decrease of several abundant 18:0 containing PE species (Figure 3B, 3C). Ether-linked 18:0 was the most abundant ether-linked hydrocarbon chain detected in our samples. This observed decrease was specific to ether-linked 18:0, as there was no statistically significant decrease in PE containing 16:0 ether-linked tails (Figure S4D).

**Figure 3:**
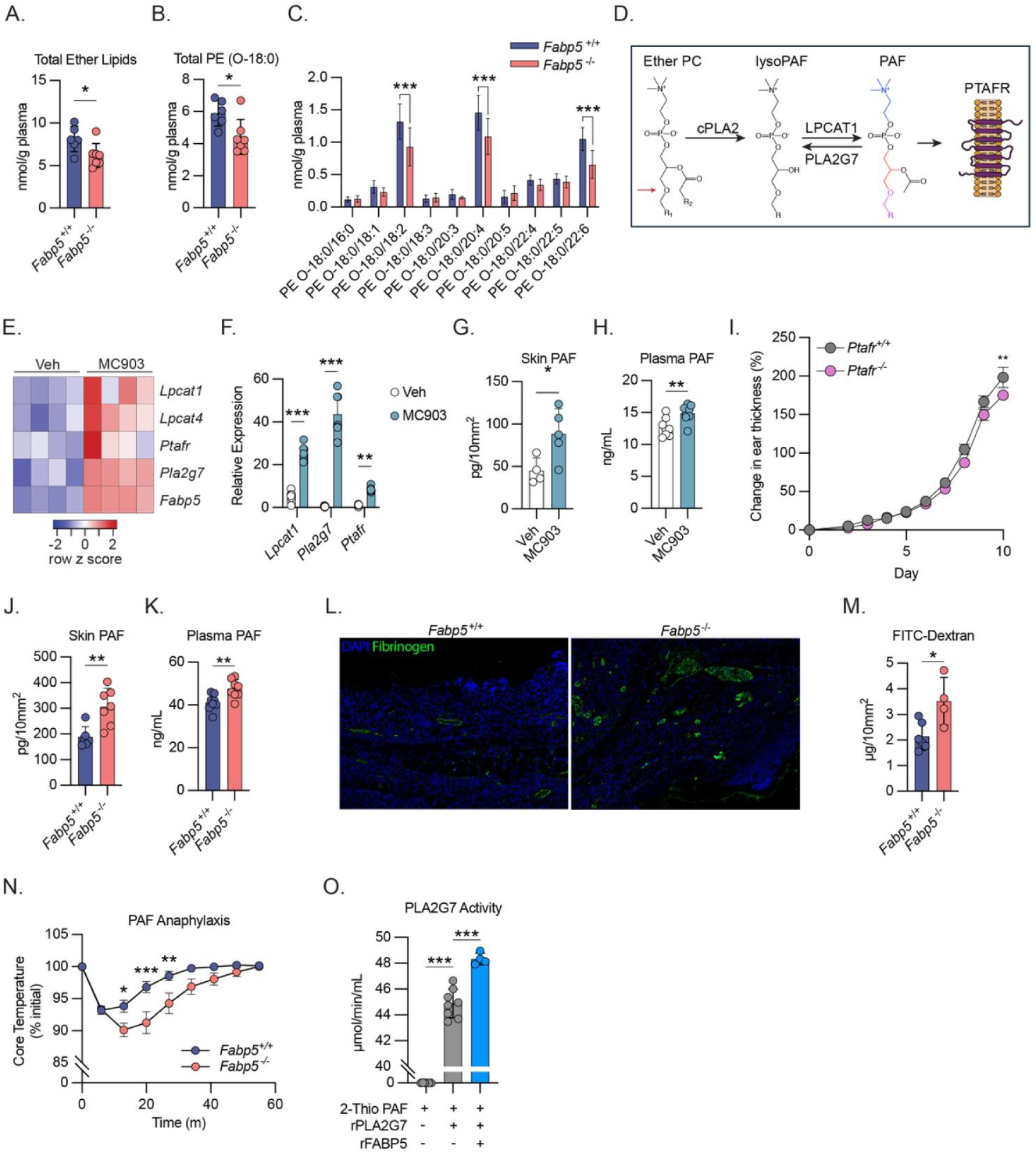
FABP5 alters ether lipid and PAF metabolism. A) High performance liquid chromatography tandem mass spectrometry (HPLC-MS/MS) based quantification of total ether-containing lipid species in plasma of *Fabp5^+/+^* and *Fabp5^-/-^* mice on day 10 of treatment with MC903. (n = 13) B) HPLC-MS/MS based quantification of total phosphatidylethanolamine (PE) species with 18:0 ether-linked tails in plasma of *Fabp5^+/+^* and *Fabp5^-/-^* mice on day 10 of treatment with MC903. (n = 13) C) HPLC-MS/MS based quantification of individual phosphatidylethanolamine species with ether-linked tails in plasma of *Fabp5^+/+^* and *Fabp5^-/-^* mice on day 10 of treatment with MC903. (n = 13). Two-way ANOVA with Šidák correction for multiple comparisons. D) Schematic of PAF synthesis via remodeling pathway. PAF synthesis begins by the hydrolysis of the sn-2 linked fatty acid tail from an ether-containing phosphatidylcholine (ether-linked tail in sn-1 position indicated by red arrow) by a phospholipase A2 thus forming an ether lyso-PC (also denoted lyso-PAF). An acetyl group is added to the sn-2 position by lysophosphatidylcholine acetyltransferase 1 (LPCAT1) forming PAF which then signals via the PAF receptor encoded by *Ptafr.* Extracellular PAF is quickly degraded by the constitutively secreted phospholipase A2, PLA2G7 (sometimes denoted PAF acetyl-hydrolase (PAF-AH)). Structural formula of PAF is colored to illustrate its metabolic components (blue: choline headgroup, red: glycerol backbone, purple: ether-linked hydrocarbon tail). E) Heat map of relative gene expression from RNAseq of naïve (vehicle treated) and dermatotic (MC903 treated) mouse skin showing expression differential expression of various genes involved in platelet activating factor (PAF) metabolism and signaling. F) qPCR analysis of *Lpcat1* and *Pla2g7* and *Ptafr* gene expression from mouse skin on day 10 of treatment with vehicle or MC903. (n = 15) G) Quantification via ELISA of PAF from mouse skin punch biopsies collected on day 10 of treatment with vehicle or MC903. PAF is normalized per unit area of skin. (n = 9) H) Quantification via ELISA of PAF in mouse plasma collected on day 10 of treatment with vehicle or MC903. (n = 16) I) Caliper measurements showing increase in ear thickness over time during MC903 treatment of *Ptafr^+/+^* and *Ptafr^-/-^* mice (n = 10). Dots show mean ± SEM. Stars represent Šidák corrected *p* values for 2-way ANOVA with multiple comparisons. J) Quantification via ELISA of PAF from skin punch biopsies from *Fabp5^+/+^* and *Fabp5^-/-^*mice on day 10 of treatment with vehicle or MC903. PAF is normalized per unit area of skin (n = 13) K) Quantification via ELISA of PAF in plasma of *Fabp5^+/+^* and *Fabp5^-/-^* mice on day 10 of treatment with vehicle or MC903. (n = 16) L) Representative immunofluorescence images of showing fibrinogen staining in skin sections from *Fabp5^+/+^* and *Fabp5^-/-^* mice on day 10 of treatment with MC903. M) Fluorescence-based quantification of FITC-Dextran in skin punch biopsies from *Fabp5^+/+^*and *Fabp5^-/-^* mice on day 10 of treatment with vehicle or MC903 10 minutes post injection with FITC-Dextran 70kD i.v.. (n = 10) N) Core body temperatures (% of initial) following PAF induced anaphylaxis in naïve *Fabp5^+/+^* and *Fabp5^-/-^* mice. (n = 10). Dots show mean ± SEM. Stars represent Šidák corrected *p* values for 2-way ANOVA with multiple comparisons. O) Cell free enzymatic activity of recombinant human PLA2G7 on 2-thio PAF with or without the addition of recombinant mouse FABP5 (400nM). (control n = 8; rFABP5 n = 4) Unless otherwise noted, all data are reported as means ± SD. * *p* < 0.05; ** *p* < 0.01, *** *p* < 0.005. Two-tailed unpaired Student’s *t*-test. Datapoints are discrete biological replicates except in (M) which displays technical replicates.

Ether lipids are a subset of glycerophospholipids which contain an ether-linked, instead of the more common acyl-linked, hydrocarbon chain at the sn-1 position of the glycerol backbone. Ether lipids have important roles in multiple cell processes including the control of membrane fluidity, cell differentiation, and oxidative stress^30^. Ether lipids are also a necessary precursor for the synthesis of platelet activating factor (PAF, 1-O-alkyl-2-acetyl-sn-glycero-3-phosphocholine), a potent bioactive lipid signaling molecule (see Figure 3D). PAF has been previously associated with allergic disease^31^ and is well-known for its functions in inducing vascular leak^31^ and as a neutrophil chemoattractant^32^. For these reasons, we decided to test whether PAF metabolism was altered in dermatitis and could be further altered by FABP5.

The biosynthesis of PAF occurs through two separate enzymatic pathways; both pathways, however, require an ether lipid precursor. Once formed, PAF signals through the PAF receptor (encoded by *Ptafr*) but is quickly deacetylated and inactivated by the A2 phospholipase, PLA2G7, which is constitutively expressed and secreted (see Figure 3D for schematic). Returning to the previously published RNAseq data from mouse skin^22^, we found that multiple genes involved in PAF metabolism were upregulated in the dermatotic skin (Figure 3E). The upregulation of these PAF-related genes was confirmed in our own MC903-treated mice (Figure 3F), suggesting that PAF metabolism may be altered during disease and contribute to disease pathology. Indeed, we found that PAF concentrations were higher in plasma and affected skin of mice treated with MC903 compared to vehicle treated controls (Figure 3G, 3H). To test the functional effects of PAF during MC903-induced dermatitis, we generated *Ptafr^-/-^* mice by excising the entire *Ptafr* coding region on mouse chromosome 4. As previously reported^33^, *Ptafr^-/-^*mice were grossly normal and displayed no obvious skin or hair defects. We treated *Ptafr^-/-^* and WT littermates with MC903 and observed a mild amelioration in dermatitis-induced ear swelling in *Ptafr^-/-^* mice, demonstrating that PAF-PTAFR signaling drives dermatitis severity in this mouse model (Figure 3I).

The observation that FABP5-deficient animals have altered concentrations of ether-linked lipids in circulation led us to reason that FABP5-deficient animals may exhibit altered PAF metabolism during dermatitis that contributes to the observed increase in disease severity. In fact, we found that MC903-treated FABP5-deficient animals had increased concentrations of PAF in plasma and skin compared to their WT counterparts (Figure 3J, 3K). Overall, these data suggest that FABP5 has broad effects on systemic ether lipid metabolism and controls the abundance of the specific bioactive ether lipid, PAF.

### FABP5 ameliorates PAF-related pathology

Among other effects, PAF is a potent modulator of endothelial cell function and induces rapid vascular leak in vivo^31^. Since MC903-treated FABP5 KO mice had increased concentrations of PAF in skin and circulation, we hypothesized that the increased dermal thickness observed in these mice may be due, in part, to an increase in vascular leak and tissue edema.

To test this hypothesis, we stained MC903-treated WT and FABP5 KO skin sections for extravascular fibrinogen, a marker of pathogenic vascular leak. We found increased extravascular fibrinogen deposited in FABP5 KO skin suggesting enhanced vascular permeability (Figure 3L). As an orthogonal approach, we i.v. injected 70kD FITC dextran into WT and FABP5 KO animals on day 10 of MC903 treatment. FITC quantification from skin punch biopsies revealed increased diffusion of FITC-dextran in FABP5 KO skin, again suggesting increased vascular permeability in these animals (Figure 3M). Together these results suggest that the increased PAF concentrations in FABP5 KO mice are associated with increased vascular leak, which may contribute to the overall increase in dermal thickness.

FABP5 could affect PAF concentrations through multiple potential mechanisms including by altering the rates of PAF synthesis or degradation. This effect could be facilitated by an ability of FABP5 to alter the abundance or location of PAF precursors, or by interacting directly with enzymes like LPCAT1 or PLA2G7 to promote PAF synthesis or degradation respectively. Intravenous administration of PAF in mice leads to an anaphylaxis-like response^34^. This response is dependent on PTAFR signaling, as PTAFR KO mice are completely protected from PAF anaphylaxis (Figure S5A). Additionally, the severity of PAF anaphylaxis is sensitive to the rate of PAF degradation, as the overexpression of PLA2G7 protects mice from PAF-induced anaphylaxis (Figure S5B, S5C). To test whether FABP5 might enhance PAF degradation in vivo, we compared the anaphylactic response to PAF in WT and FABP5 KO mice. FABP5 KO mice were more susceptible to PAF-induced anaphylaxis suggesting FABP5 might contribute to PAF degradation in vivo (Figure 3N). To further test this hypothesis, we used a cell free assay that measures the enzymatic activity of PLA2G7 on the deacetylation of 2-thio PAF. Addition of recombinant mouse FABP5 to this in vitro reaction, increased PLA2G7 activity (Figure 3O). These results suggest that FABP5 may alter PAF metabolism by enhancing PAF degradation by PLA2G7.

### Modulating PAF metabolism alters dermatitis severity dependent on FABP5

We next sought to test whether pharmacologically altering PAF metabolism in vivo could recapitulate the effects of FABP5 deficiency. We gavaged WT and FABP5-deficient mice with the well-characterized PLA2G7 inhibitor (PLA2G7i), darapladib, during the course of MC903 treatment (Figure 4A). We hypothesized that inhibiting the major degrative enzyme of PAF would exacerbate dermatitis in WT mice and recapitulate the phenotype observed in FABP5 KO animals. Indeed, WT mice treated with daily PLA2G7i displayed increased skin thickness compared to their vehicle treated counterparts (Figure 4B). This exacerbated skin thickness mirrored the disease severity of vehicle treated *Fabp5^-/-^* animals (Figure 4B). Surprisingly, we observed no further exacerbation of skin thickness in *Fabp5^-/-^*animals treated with PLA2G7i, which may indicate that PAF signaling in these animals may already be at the upper limit of its relevant physiological range (Figure 4B). Congruent with these changes in skin thickness, we detected increased extravasated FITC dextran in PLA2G7i-treated WT skin (Figure 4C). Finally consistent with the role of PAF as a neutrophil chemoattractant, we found that PLA2G7i treatment increased neutrophil infiltration into WT skin but again had no effect in *Fabp5^-/-^*mice (Figure 4D).

**Figure 4:**
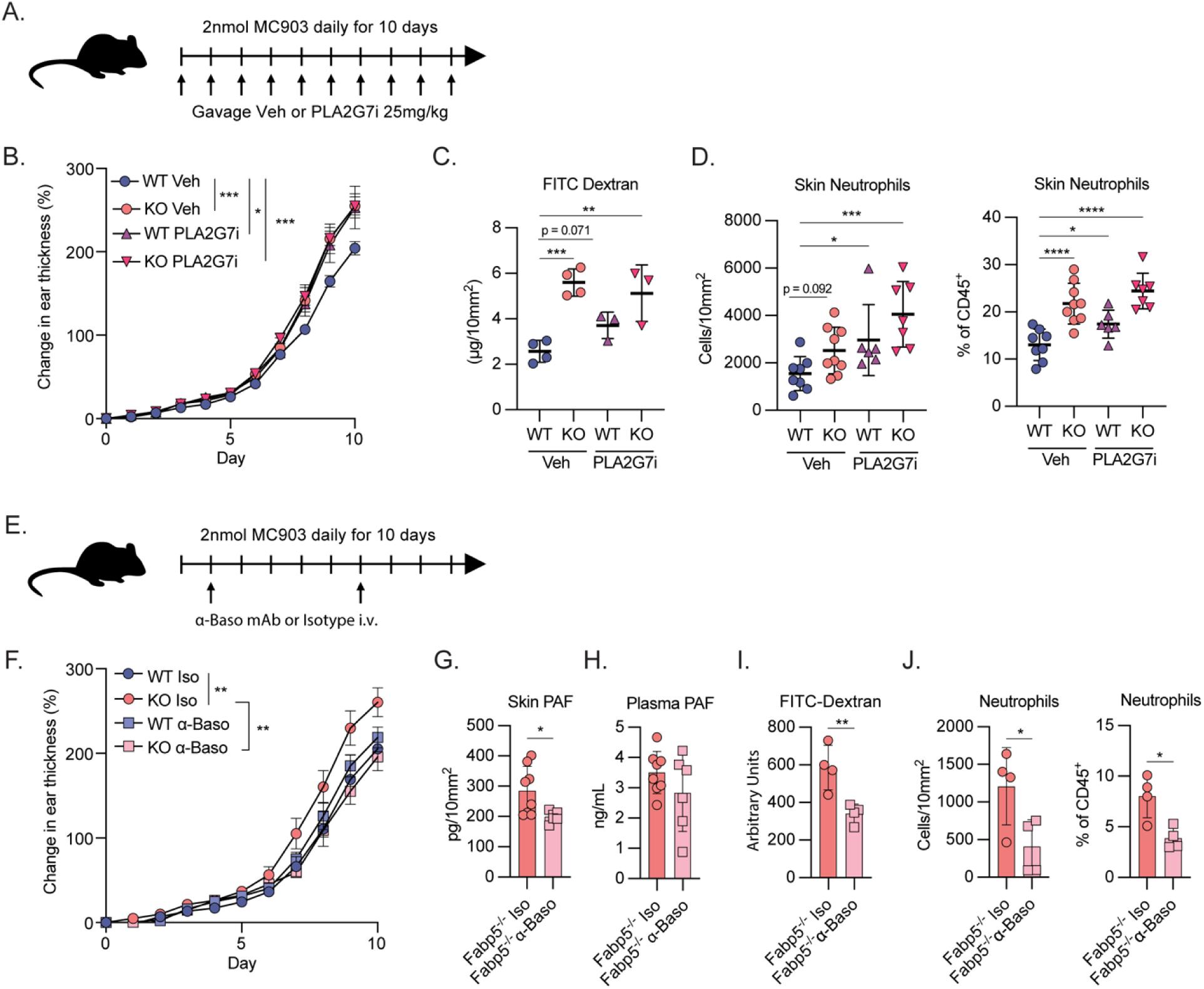
Modulating PAF metabolism alters dermatitis severity dependent on FABP5. A) Schematic of experimental design for inhibition of PLA2G7 during MC903-induced dermatitis. B) Caliper measurements showing increase in ear thickness over time during MC903 treatment of *Fabp5^+/+^*(WT) and *Fabp5^-/-^* (KO) mice gavaged with daily vehicle or PLA2G7i (n = 16). Dots show mean ± SEM. Stars represent day 10 Šidák corrected *p* values for 2-way ANOVA with multiple comparisons. C) Fluorescence-based quantification of FITC-Dextran in skin punch biopsies from MC903-treated *Fabp5^+/+^* (WT) and *Fabp5^-/-^* (KO) mice gavaged with daily vehicle or PLA2G7i on day 10 of treatment. (n = 14) D) Flow cytometry-based quantification of total neutrophils in skin of MC903-treated *Fabp5^+/+^*(WT) and *Fabp5^-/-^* (KO) mice gavaged with daily vehicle or PLA2G7i on day 10 of treatment. Cell count is normalized per unit area (left) or reported as a percent of total CD45^+^ immune cells (right). (n = 30) E) Schematic of experimental design for depletion of basophils during MC903-induced dermatitis. F) Caliper measurements showing increase in ear thickness over time during MC903 treatment of *Fabp5^+/+^*(WT) and *Fabp5^-/-^* (KO) mice treated with basophil depleting antibody or isotype control. (n = 13). Dots show mean ± SEM. Stars represent day 10 Šidák corrected *p* values for 2-way ANOVA with multiple comparisons. G) Quantification of PAF in skin punch biopsies on day 10 from MC903-treated *Fabp5^-/-^* mice treated with basophil depleting antibody or isotype control. PAF is normalized per unit area of skin (n = 14) H) Quantification of PAF in plasma on day 10 from MC903-treated *Fabp5^-/-^* mice treated with basophil depleting antibody or isotype control. (n = 14) I) Fluorescence-based quantification of FITC-Dextran in skin punch biopsies on day 10 from MC903-treated *Fabp5^-/-^* mice treated with basophil depleting antibody or isotype control. Reported in arbitrary fluorescent units. (n = 8) J) Flow cytometry-based quantification of total neutrophils in skin on day 10 from MC903-treated *Fabp5^-/-^* mice treated with basophil depleting antibody or isotype control. Cell count is normalized per unit area (left) or reported as a percent of total CD45^+^ immune cells (right). (n = 8) Unless otherwise noted, all data are reported as means ± SD. * *p* < 0.05; ** *p* < 0.01, *** *p* < 0.005. Two-tailed unpaired Student’s *t*-test. Datapoints are discrete biological replicates. (C) and (D) were analyzed by one-way ANOVA with Fisher’s least significant difference test.

There are no pharmacologic reagents to block PAF synthesis, which occurs through multiple pathways. However, basophils are a major cellular source of PAF^34^ and are highly prevalent in the dermatotic skin following MC903 treatment^35^. We therefore hypothesized that depleting basophils may reduce the elevated PAF concentrations found in *Fabp5^-/-^* mice and ameliorate disease. We depleted basophils in MC903-treated *Fabp5^+/+^* and *Fabp5^-/-^* animals by i.v. administration of an anti-CD200R3 antibody, which results in the systemic depletion of both basophils and mast cells (Figure 4E). Basophil depletion protected *Fabp5^-/-^* mice from the exacerbated skin thickening observed in their isotype treated counterparts. (Figure 4F). Basophil depletion also lowered PAF concentrations in the skin, although plasma concentrations were not affected (Figure 4G, 4H). Additionally, consistent with the role of PAF in promoting vascular permeability and neutrophil infiltration, we found decreased tissue FITC-dextran (Figure 4I) and fewer neutrophils (Figure 4J) in the skin of basophil-depleted FABP5 KO mice. Altogether these results suggest that regulation of PAF metabolism through an axis dependent on FABP5 and basophils controls alters dermatitis severity.

## Discussion

The elevated expression of FABP5 in the skin of patients with AD was documented more than a decade ago^36^, yet the functional role of this protein during disease has remained enigmatic. In this study, we characterize FABP5 expression in a mouse model of AD and demonstrate that FABP5 upregulation in skin and circulation during allergic skin inflammation is conserved in mice (Figure 1A-D) as previously reported in humans^21^. We find that FABP5 expression does not exacerbate disease, as hypothesized by previous in vitro work^21^, but rather is an important factor that constrains pathology (Figure 1E, 1F), and therefore its upregulation represents a negative feedback loop aimed at dampening allergic skin inflammation. This conclusion supports previous work that has demonstrated FABP5 to be a protective factor in mouse models of allergic airway inflammation, suggesting that FABP5 may be playing a fundamental role in the regulation of type 2 immunity.

We next used conditional genetics to determine the cell types in which FABP5 expression was operative to constrain disease but found that deleting FABP5 from neither keratinocytes nor myeloid cells could recapitulate the exacerbated disease observed in the germline FABP5 KO mice (Figure 2H, 2I). This result points to several possible mechanistic roles for FABP5. First, FABP5 may be exerting its function through a cell type outside of the keratinocyte or myeloid lineages. While we explored in depth the expression patterns of FABP5 in the cells resident in the skin, FABP5 is expressed widely throughout the body^15^, and expression in other cell types, such as neurons, hepatocytes, or endothelial cells, may be crucial to control disease. Another possibility is that FABP5 is playing a role in lipid metabolism that is shared and redundant among cell types. Finally, it’s possible that extracellular FABP5, which is derived from multiple cell types, is crucial for constraining disease. Since circulating FABP5 was observed in both *K14ΔFabp5* and *LysMΔFabp5* strains (Figure 2K, 2M), we were unable to parse potential intracellular and extracellular functions of the protein. Further study and mechanistic insight will be necessary to test these possibilities.

Notably, we uncovered a dramatic and specific alteration in systemic lipid metabolism in FABP5-deficient mice. Specifically, FABP5-deficiency resulted in a dramatic decrease in the abundance of ether-linked glycerophospholipids in plasma (Figure 3A-C). To our knowledge, this is the first report describing a role for FABP5 in directing systemic lipid metabolism. Among other functions, ether-linked lipids are a necessary precursor of PAF, and we observed increased PAF concentrations in FABP5-deficient animals with dermatitis (Figure 3J, 3K). The mechanism by which FABP5 modulates PAF concentration during dermatitis remains uncertain. However in vitro and in vivo experiments suggest FABP5 could enhance the PLA2G7-mediated degradation of PAF to lyso-PAF (Figure 3N, 3O). Whether this function of FABP5 effects the abundance of systemic ether lipids, or whether effects on general ether lipid metabolism are the result of a distinct function of FABP5 is unclear.

PAF is a potent inducer of vascular leak^31^ and a neutrophil chemoattractant^32^. Indicative of these functions, we found that dermatitis in FABP5-deficient animals was associated with increased endothelial leak, and neutrophils in the skin (Figure 3L, 3M; Figure 1J). We therefore hypothesize that increased PAF concentrations in FABP5-deficient mice are responsible for this specific phenotype of exacerbated disease. Supporting this supposition, inhibition of the PAF degradative enzyme, PLA2G7, exacerbated dermatitis, vascular leak and neutrophil infiltration while depletion of basophils, a major cellular source of PAF, ameliorated disease (Figure 4). FABP5 has previously been demonstrated to be protective in multiple mouse models of inflammation including allergic airway inflammation^19,20^ and sepsis^37^. The effect of FABP5 on systemic lipid metabolism was not assessed in these studies, however as PAF has been implicated as an inflammatory mediator in both allergic airway inflammation^38^ and sepsis^39^, it is tempting to speculate that at least part of the protective effects of FABP5 in these models could be explained by its modulation of PAF and ether-lipid metabolism. Additional research will be required to parse out intracellular and systemic effects of FABP5 deficiency in these models and to explore whether the effect of FABP5 on PAF and ether lipid metabolism could be a conserved phenomenon across various types of inflammation.

PAF was discovered in the 1970s as a basophil derived mediator of platelet aggregation and secretion of histamine^31^. Since then, much data has been generated implicating PAF and its degradative enzyme, PLA2G7 (sometimes denoted as platelet activating factor acetylhydrolase or PAF-AH), in the pathology of several diseases, including asthma^38,40^, anaphylaxis^41,39^, and sepsis^39^. The role of PAF in the skin is less clear. While intradermal injection of PAF induces a local biphasic reaction^42^, the development of PAF inhibitors for the treatment of AD has not yet been successful. Still, several studies have explored the roles PAF and PAF metabolism in the skin. A recent study examined the transcriptional profiles of skin from patients with spontaneously resolved AD using scRNAseq and found that PLA2G7 was upregulated in melanocytes of patients with spontaneously healed AD compared to healthy controls or patients with active AD^43^. These intriguing results are in concordance with our conclusions and suggest that the story of PAF and PLA2G7 in the skin is still unfolding and warrants further research.

## Materials and Methods

### Mouse strains

*Fabp5^-/-^*, *Fabp5^HA^*, and *Fabp5^flox/flox^* and *Ptafr^-/-^* mice were generated in our in-house CRISPR Core and are available upon request. *Rag2^-/-^Il2rg^-/y^*DKO mice were purchased from Taconic (4111-M) and crossed with *Fabp5^-/-^* mice to obtain *Rag2^-/-^Fabp5^-/-^* DKO and *Rag2^-/-^* controls. *K14-cre* (JAX 018964) and *LysM-cre* (JAX 004781) mice were purchased from Jackson Laboratory and crossed with *Fabp5^flox/flox^* mice to generate conditional KO strains. For all experiments, mice were aged 8-12 weeks in cohoused and sex-matched conditions to minimize microbial variation. Mice were housed with 14:10 h day:night cycle and were fed a standard chow. Both male and female mice were used in experiments. All mice were bred and maintained in the facilities of the Yale Animal Resources Center and all animal experimentation was performed in compliance with Yale Institutional Animal Care and Use Committee protocols.

### MC903 treatment, inhibitor treatment, and basophil depletion

For MC903 treatment, mice were anesthetized with isofluorane before application 2nmol of MC903 (Calcipitriol, Tocris Cat. No. 2700) in 20uL of ethanol onto skin (one or both pinnae, or shaved dorsal skin as indicated). 20uL of ethanol only was used as vehicle only control where indicated. MC903 or vehicle was applied each day for 10 days. Skin thickness was measured daily via electronic calipers (Mitutoyo PK-0505CPX). In experiments with PLA2G7i, mice of each genotype were randomized into PLA2G7i or vehicle control groups and gavaged daily with PLA2G7i, darapladib, (25mg/kg Sigma, SML3097) or vehicle control (100uL H_2_0 + 5% DMSO) coincident with MC903 treatment. In experiments with basophil depletion, mice of each genotype were randomized into α-Baso or isotype control groups on day 1 of MC903 treatment. Mice were injected i.v. with 30ug of a basophil depleting antibody (Biologend UltraLEAF clone Ba160) or equal amount of Rat IgG2b isotype control (Biolegend 400602) on treatment days 1 and 6.

### FITC-Dextran injections and analysis

On day 10 of MC903 treatment mice were injected i.v. with 200uL of 50mg/mL FITC Dextran 70kD (Sigma 46945) dissolved in PBS. Mice were sacrificed 10 minutes after injection and punch biopsies (2 x 2mm or 6mm) were taken from ear pinnae and flash frozen. For analysis, tissue biopsies were homogenized in 1mL ice cold PBS using a Kinematica Polytron® PT2100 tissue homogenizer. Homogenates were spun at 10 000 x g for 10 min at 4C and liquid fraction was collected, transferred to a black 96 well assay plate, and FITC fluorescence was measured using a BioTek Synergy HTX (Excitation:485/20 Emission:528/20). Fluorescent values were transformed to μg/mL using a standard curve of FITC Dextran 70kD diluted in PBS at known concentrations and then normalized to punch biopsy area. In some experiments fluorescent values are reported untransformed as arbitrary fluorescent units.

### PAF Induced Anaphylaxis

Core body temperature of mice was obtained via rectal probe (Physitemp; TH-5 Thermalert) before i.v. injection of 500ng PAF C16 (Sigma 511075) in 200μL of PBS. Core body temperature was measured every 6 minutes for 60-80 minutes.

### Hydrodynamic Injection

Mouse weights were recorded and mice were injected via tail vain with a saline solution corresponding 10% of body weight with or without 25μg of mouse Pla2g7 plasmid (Origene NM_013737).

### Gene Expression Analysis

Mice were sacrificed, and skin punch biopsies were collected and immediately submerged in RNA*later*™ (Invitrogen AM7021). Biopsies were stored at 4C until RNA could be extracted. Biopsies were later homogenized in 1mL ice cold Trizol using a Kinematica Polytron® PT2100 tissue homogenizer, and whole RNA was extracted. RNA concentrations were normalized, and cDNA was synthesized using Maxima H Minus Reverse Transcriptase (ThermoFisher EP0753). qPCR was performed with BioRad C1000 Touch Thermocycler with CFX384 Real Time System using iTaq Universal SYBR Green Super Mix (BioRad 1725124) and 0.5 μmol/L primers. Relative expression values were normalized to control gene (rRNA 36B4) and expressed in terms of linear relative mRNA values. Primers for Lpcat1 and Pla2g7 were obtained from PrimerBank^44–46^ (PrimerBankID 148747362c3, 31980752a1). The sequences for other primers can be found below.

Fabp4 Forward: GGATGGAAAGTCGACCACAA; Reverse: TGGAAGTCACGCCTTTCATA
Fabp5 Forward: ACGGCTTTGAGGAGTACATGA; Reverse: CTCGGTTTTGACCGTGATG
Ptafr Forward: CAGGCCACAACACAGAGGCTCG; Reverse: TCACCTGGTCATGGAGCGCTGA
36b4 Forward: AGATGCAGCAGATCCGCAT; Reverse: GTTCTTGCCCATCAGCACC

### Preparation of serum and skin homogenates, and ELISAs

#### Serum

mouse blood was collected via retroorbital bleed and allowed to clot at room temperature for 30 min. Clotted blood was spun at 2000 x g for 10 min, and serum was collected and frozen until analysis.

#### Skin

punch biopsies (3mm or 4mm) were collected and snap frozen in liquid N_2_. Punch biopsies were later homogenized in 1mL ice cold PBS using a Kinematica Polytron® PT2100 tissue homogenizer. Homogenates were spun at 10 000 x g for 10 min at 4C and liquid fraction was collected for downstream analysis.

#### ELISAs

FABP5 ELISA (MBL; CY-8056) or PAF ELISA (Abcam; ab287801) were used on mouse serum or mouse skin homogenates according to manufacturer’s instruction and resulting calculated concentrations were normalized to punch biopsy area.

### PLA2G7 Activity Assay

PLA2G7 enzymatic deacetylation of 2-thio PAF was measured from murine serum using a PAF Acetylhydrolase Assay Kit (Cayman 760901) according to manufacturer’s instructions. In cell free assays, the effect of recombinant murine FABP5 (Cayman 10007433) on recombinant human PLA2G7 deacetylation of 2-thio PAF was measured via PAF Acetylhydrolase Inhibitor Screening Assay Kit (Cayman 10004380) according to manufacturers instructions.

### H&E histology and histological measurements

Mice were euthanized, and ear pinnae were excised, sandwiched between blotting paper and fixed in 10% neutral buffered formalin overnight. Pinnae were bisected and embedded in paraffin, transverse cuts were stained with hematoxylin and eosin by Yale Pathology Tissue Services. Brightfield images of H&E slides were captured using a Keyence BZ-X810 microscope using the 10x objective, and images were stitched together to recreate whole tissue sections for histological measurements. Dermal and epidermal thickness was measured 4mm from apex of pinnae using ImageJ.

### Immunofluorescence staining and visualization

For FABP5-HA staining in mouse skin, shaved dorsal skin was treated for 10 days with vehicle or MC903 before being excised and fixed in 4% paraformaldehyde for 4 hours at 4C. Fixed tissue was dehydrated in 30% sucrose and then embedded in TissueTek O.C.T. compound for sectioning. 10μm cryostat-cut, transverse skin sections were washed, permeabilized in PBS 0.3% Triton X-100, blocked for 1hr with mouse IgG_1_ isotype control, and incubated with chicken α-Keratin 14 1:200 (Biolegend; 906004) rabbit α-Keratin 10 1:200 (Biolegend; 905403) and mouse α-ΗΑ AF647 1:200 (Biolegend; 682404) followed by goat α-chicken IgY AF568 1:1000, donkey α-rabbit IgG AF488 1:1000 secondary antibodies and DAPI counterstain. For FABP5-HA staining in skin-resident immune cells, CD45^+^ cells were isolated from mouse skin via FACS before being adhered to slides via Cytospin. Cells were then fixed, permeabilized and stained with rabbit α-ΗΑ 1:1000 followed by donkey α-rabbit IgG AF488 secondary and DAPI counterstain. For fibrinogen staining, formalin fixed, paraffin embedded tissues were cut and rehydrated. Antigen retrieval was achieved by boiling slides in 10mM citrate buffer pH6 for 20 min. Tissue sections were permeabilized as above before being blocked for 1 h with 5%BSA in PBS 0.1% Triton X-100. Tissue sections were then incubated in α-human fibrinogen 1:100 (Agilent Technologies; A008002-2) followed by donkey α-Rabbit AF488 1:1000 secondary and DAPI counterstain. In all cases, immunofluorescence images were captured using a Keyence BZ-X810 epifluorescent microscope using 10x, 20x, and 40x objectives.

### Isolation of immune cells from mouse skin

Mice were euthanized and 5mm or 6mm punch biopsies were taken from ear pinnae and plunged in ice cold PBS. Ventral and dorsal dermal sheets were separated, and biopsies were minced and placed in digestion buffer (RPMI, 0.25mg/mL Liberase TM, 0.5mg/mL DNase). Minced biopsies were digested for 45 min at 37C with gentle shaking at 180RPM. Following digestion, the suspension was passed through a 70um cell strainer with the aid of a 3mL syringe plunger. Cells were washed twice (RPMI 5% FBS) before immunocytes were enriched via a single 35% Percoll gradient.

### Flow cytometric analysis

Isolated immune cells were stained with appropriate antibody cocktail and fixable viability dye diluted into FACS buffer (PBS 2% FBS) for 20min at 4C. Cells were washed and fixed with BD Cytofix/Cytoperm (BD 554714) for later analysis. For intracellular cytokine staining, isolated immune cells were first resuspended in RMPI 10% FBS and stimulated with PMA/Ionomycin for 3h at 37C. BD Golgi Plug™ (BD 555029) was added (1:1000) for the last 90 min. Cells were then washed and stained as above before being fixed and permeabilized, for intracellular cytokine staining overnight at 4C. For intracellular FABP5-HA staining, isolated cells were stained with antibody cocktail for appropriate surface antigens before being fixed and permeabilized using the eBioscience™ Foxp3/Transcription Factor Fixation/Permeabilization kit. Fixed cells were stained for intracellular HA overnight at 4C. All antibody clones, catalog numbers and dilutions used for staining can be found in Supplementary Table 1. AccuCheck cell counting beads (Invitrogen PCB100) were added to all samples for cell number quantification. Calculated cell numbers were then normalized to punch biopsy area. All flow cytometry data was collected on BD LSR2 with FACSDiva 7 software. Flow cytometry data were analyzed with FlowJo (v10.10.0).

### scRNA-seq sample preparation

Five RAG2 KO mice were treated with MC903 and vehicle on contralateral ears for 10 days, and ear thickness was measured each day using calipers. All 5 vehicle treated ears and 3 MC903 treated ears were pooled respectively and skin was digested to isolate immunocytes. Cells were stained, and two populations, innate lymphocytes (Live CD45+Ly6G-Lin-CD90.2+) and myeloid (Live CD45+Ly6G-CD90.2-) were isolated via FACS. Isolated cells were then recombined for a final ratio of 30% innate lymphocytes and 70% myeloid.

scRNA-seq libraries from the recombined cell populations were prepared Chromium Next GEM Single Cell 3′ v3.1 Library & Gel Bead Kit (10X Genomics) according to the manufacturer’s instructions. Briefly, emulsions were generated using the 10X Chromium Controller for a targeted recovery of 10,000 cells. The barcoded cDNAs were isolated from the emulsion and amplified by PCR (12 cycles). The amplified cDNAs were subjected to fragmentation, end repair, and A-tailing, before sample indexing by PCR (16 cycles). The resulting libraries were sequenced by NovaSeq 6000.

### scRNA-seq data analysis

Raw sequencing data were processed using CellRanger software (v7.1.0) and aligned to the mouse mm10-2020-A reference genome. The data were then filtered by removing genes expressed in fewer than 3 cells and cells with fewer than 200 detected features. Further analysis was performed using Seurat R package (v5.0.1). First, samples were combined, and poor-quality cells were identified as cells with number of expressed genes <200 or >7,500, or mitochondrial gene percentage over 5%, and excluded. Samples were normalized via SCTransform and principal components were identified via RunPCA. The first 30 principal components were used for cell clustering. FindAllMarkers was then used to identify cluster specific genes which were used to manually annotate cell clusters. Multiple fibroblast and endothelial clusters were respectively combined for ease of visualization.

### Lipidomics analysis

EDTA plasma was collected from mice on day 10 of MC903 treatment and immediately frozen until analysis. Quantitative untargeted lipidomic analysis on plasma samples was performed using C18 HPLC-MS/MS by the Northwest Metabolomics Research Center at the University of Washington.

### Statistical Analysis

All data were graphed and statistical analyses performed using Graphpad Prism 10 (v10.4.0). Unless otherwise indicated all data is plotted as mean ± SD. Statistical significance was determined via two-tailed unpaired Student’s *t* test, one-way ANOVA, or two-way ANOVA with corrections for multiple comparisons as indicated in figure legends. *p* values < 0.05 were considered statistically significant.

## Data Availability

Source data from scRNA seq will be made publicly available before publication.

## Acknowledgments

We would like to thank J. Alderman, C. Hughes, L. Evangelisti, E. Hughes-Picard, Bill Philbrick, and J. Harrocks for administrative and technical assistance. We thank J.R. Brewer, M. Oh, E. Han, I. Odell, S. F. Lau and all members of the Flavell lab for thoughtful discussion and feedback throughout this project. We thank Yale Flow Cytometry for their assistance with analyzers and cell sorting. The Core is supported in part by an NCI Cancer Center Support Grant # NIH P30 CA016359. Finally, we thank Kevin Williams, the UCLA Lipidomics core and the Northwest Metabolomics Research Center at the University of Washington for their technical assistance and support in the design and execution of our lipidomics experiments. M.H.S. is supported by the NSF GRFP and the Yale Trudeau Fellowship. W.K.M. is supported by NIH T32-DK007356 and is the recipient of a Cancer Research Institute/Irvington Postdoctoral Fellowship.

## Author Contributions

M.H.S. conceived the project, designed and performed experiments, analyzed the data, and wrote the manuscript. W.K.M. assisted with experimental design, performed scRNAseq sample preparation and analyzed bulk RNAseq data. H.N.B. assisted with in vivo experiments. H.M. assisted in histological analysis and interpretation. F.Z. performed the scRNAseq alignment and assisted with analysis. A.G.Y. and R.A.F. provided resources and funding, contributed ideas to the development of this project, and supervised the drafting of the manuscript.

---

**Supplementary Table 1.**
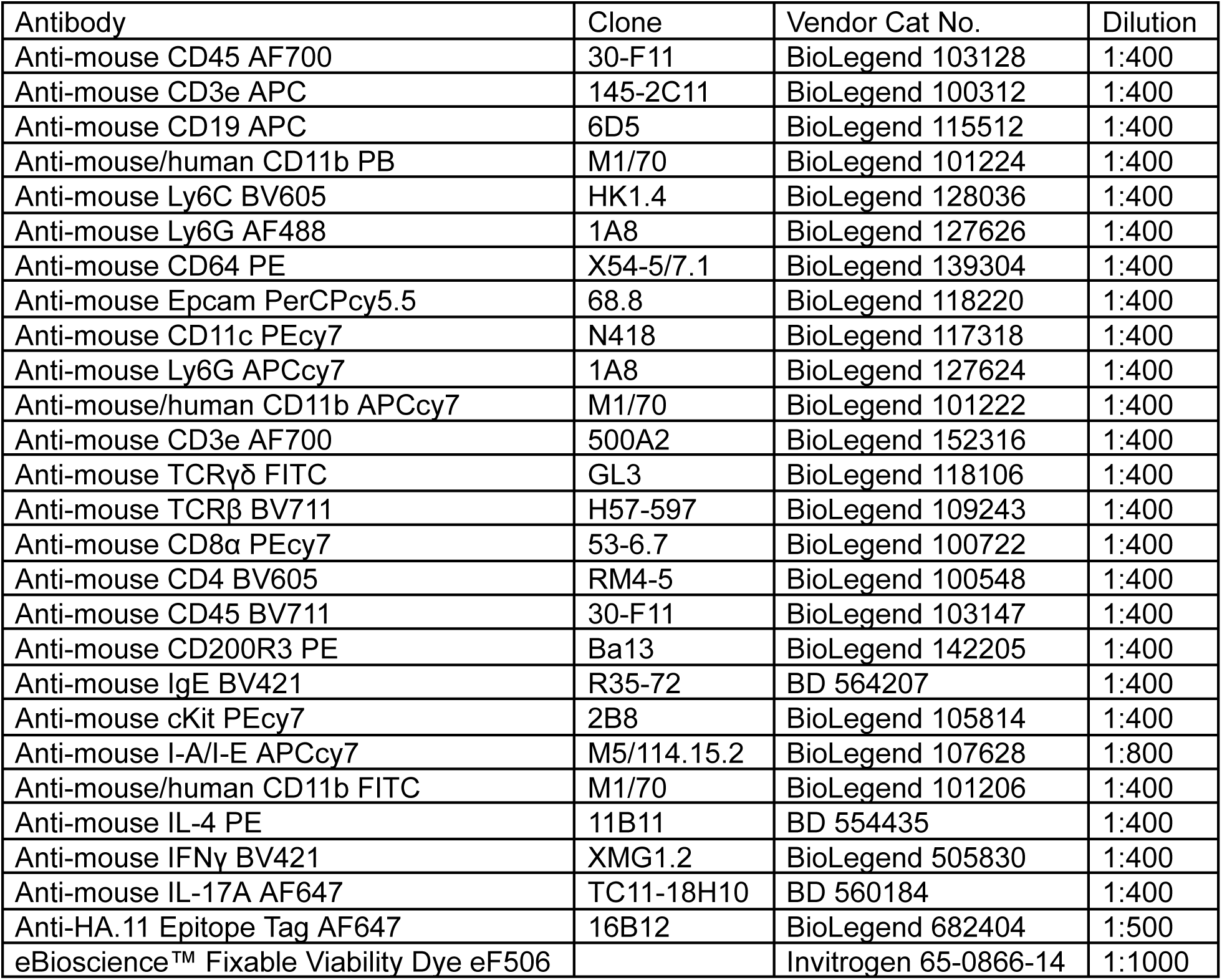
List of antibodies and their working dilutions for use in flow cytometry analysis.

| Antibody | Clone | Vendor Cat No. | Dilution |
| --- | --- | --- | --- |
| Anti-mouse CD45 AF700 | 30-F11 | BioLegend 103128 | 1:400 |
| Anti-mouse CD3e APC | 145-2C11 | BioLegend 100312 | 1:400 |
| Anti-mouse CD19 APC | 6D5 | BioLegend 115512 | 1:400 |
| Anti-mouse/human CD11b PB | M1/70 | BioLegend 101224 | 1:400 |
| Anti-mouse Ly6C BV605 | HK1.4 | BioLegend 128036 | 1:400 |
| Anti-mouse Ly6G AF488 | 1A8 | BioLegend 127626 | 1:400 |
| Anti-mouse CD64 PE | X54-5/7.1 | BioLegend 139304 | 1:400 |
| Anti-mouse Epcam PerCPcy5.5 | 68.8 | BioLegend 118220 | 1:400 |
| Anti-mouse CD11c PEcy7 | N418 | BioLegend 117318 | 1:400 |
| Anti-mouse Ly6G APCcy7 | 1A8 | BioLegend 127624 | 1:400 |
| Anti-mouse/human CD11b APCcy7 | M1/70 | BioLegend 101222 | 1:400 |
| Anti-mouse CD3e AF700 | 500A2 | BioLegend 152316 | 1:400 |
| Anti-mouse TCR$\gamma\delta$ FITC | GL3 | BioLegend 118106 | 1:400 |
| Anti-mouse TCRβ BV711 | H57-597 | BioLegend 109243 | 1:400 |
| Anti-mouse CD8α PEcy7 | 53-6.7 | BioLegend 100722 | 1:400 |
| Anti-mouse CD4 BV605 | RM4-5 | BioLegend 100548 | 1:400 |
| Anti-mouse CD45 BV711 | 30-F11 | BioLegend 103147 | 1:400 |
| Anti-mouse CD200R3 PE | Ba13 | BioLegend 142205 | 1:400 |
| Anti-mouse IgE BV421 | R35-72 | BD 564207 | 1:400 |
| Anti-mouse cKit PEcy7 | 2B8 | BioLegend 105814 | 1:400 |
| Anti-mouse I-A/I-E APCcy7 | M5/114.15.2 | BioLegend 107628 | 1:800 |
| Anti-mouse/human CD11b FITC | M1/70 | BioLegend 101206 | 1:400 |
| Anti-mouse IL-4 PE | 11B11 | BD 554435 | 1:400 |
| Anti-mouse IFNγ BV421 | XMG1.2 | BioLegend 505830 | 1:400 |
| Anti-mouse IL-17A AF647 | TC11-18H10 | BD 560184 | 1:400 |
| Anti-HA.11 Epitope Tag AF647 | 16B12 | BioLegend 682404 | 1:500 |
| eBioscience™ Fixable Viability Dye eF506 |  | Invitrogen 65-0866-14 | 1:1000 |

**Supplemental Figure 1.**
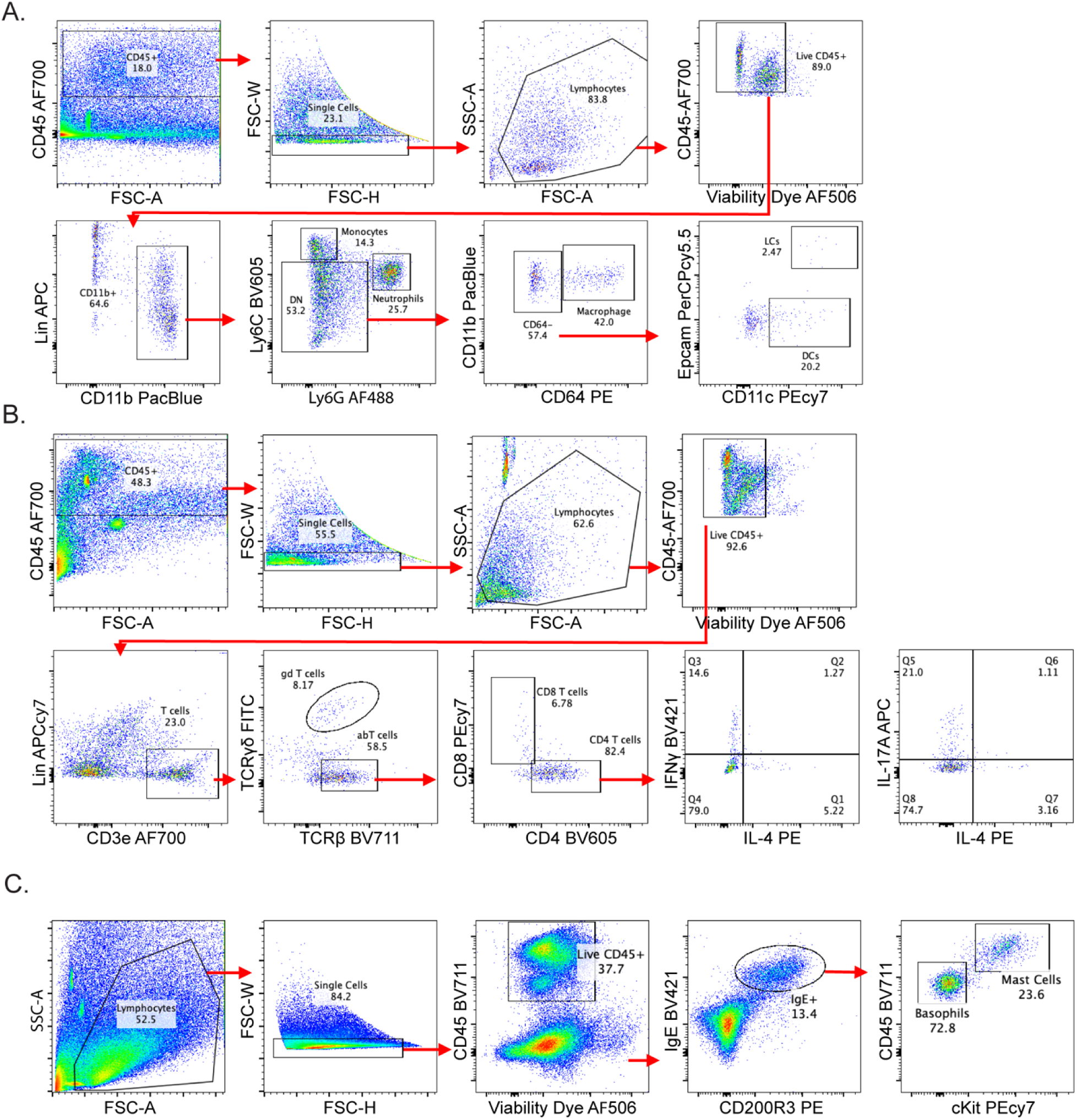
A) Flow cytometry gating strategy for quantification of total CD45+ immune cells, monocytes, neutrophils, macrophages, dendritic cells (DCs) and Langerhans cells (LCs) isolated from the skin of MC903-treated mice. B) Flow cytometry gating strategy for quantification of CD4^+^ and CD8^+^ T cells as well as the different cytokine secreting CD4+ T cell populations from the skin of MC903-treated mice. C) Flow cytometry gating strategy for quantification of basophils and mast cells from the skin of MC903-treated mice.

**Supplemental Figure 2.**
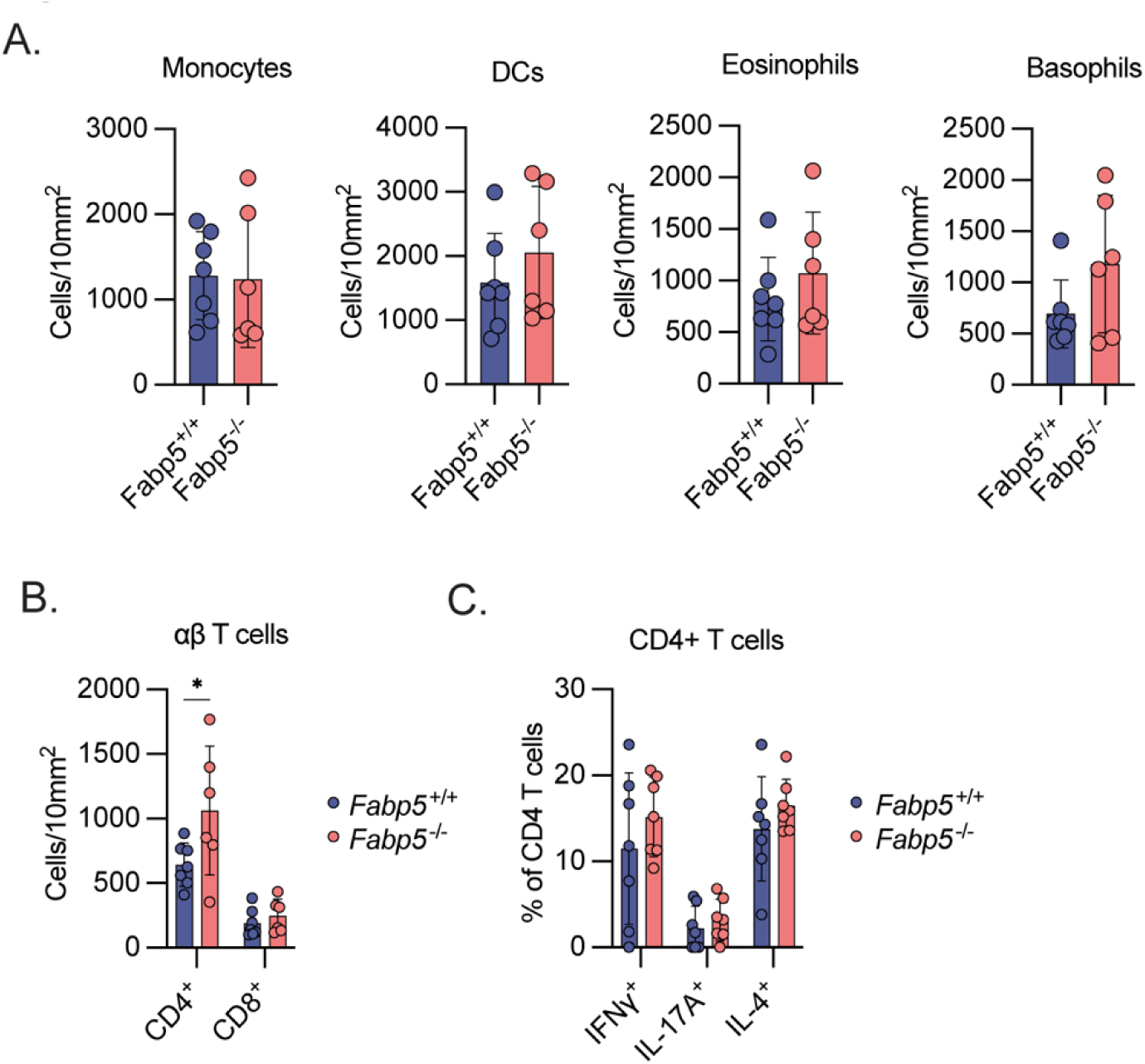
A) Flow cytometry-based quantification of total monocytes, dendritic cells (DCs), eosinophils, and basophils in skin of *Fabp5^+/+^* and *Fabp5^-/-^* mice on day 10 of treatment with MC903. Cell count is normalized per unit area. (n = 13) B) Flow cytometry-based quantification of CD4+ and CD8+ αβ T cells in skin of *Fabp5^+/+^* and *Fabp5^-/-^* mice on day 10 of treatment with MC903. Cell count is normalized per unit area. (n = 13) C) Flow cytometry-based quantification of intracellular cytokine staining of CD4^+^ T cells in skin of *Fabp5^+/+^* and *Fabp5^-/-^* mice on day 10 of treatment with MC903. Cell count is normalized as percent of total CD4^+^ T cells (n = 14) Unless otherwise noted, all data are reported as means ± SD. * *p* < 0.05; ** *p* < 0.01, *** *p* < 0.005. Two-tailed unpaired Student’s *t*-test. Datapoints are discrete biological replicates. (C) and (D) were analyzed by one-way ANOVA with Fisher’s least significant difference test.

**Supplemental Figure 3:**
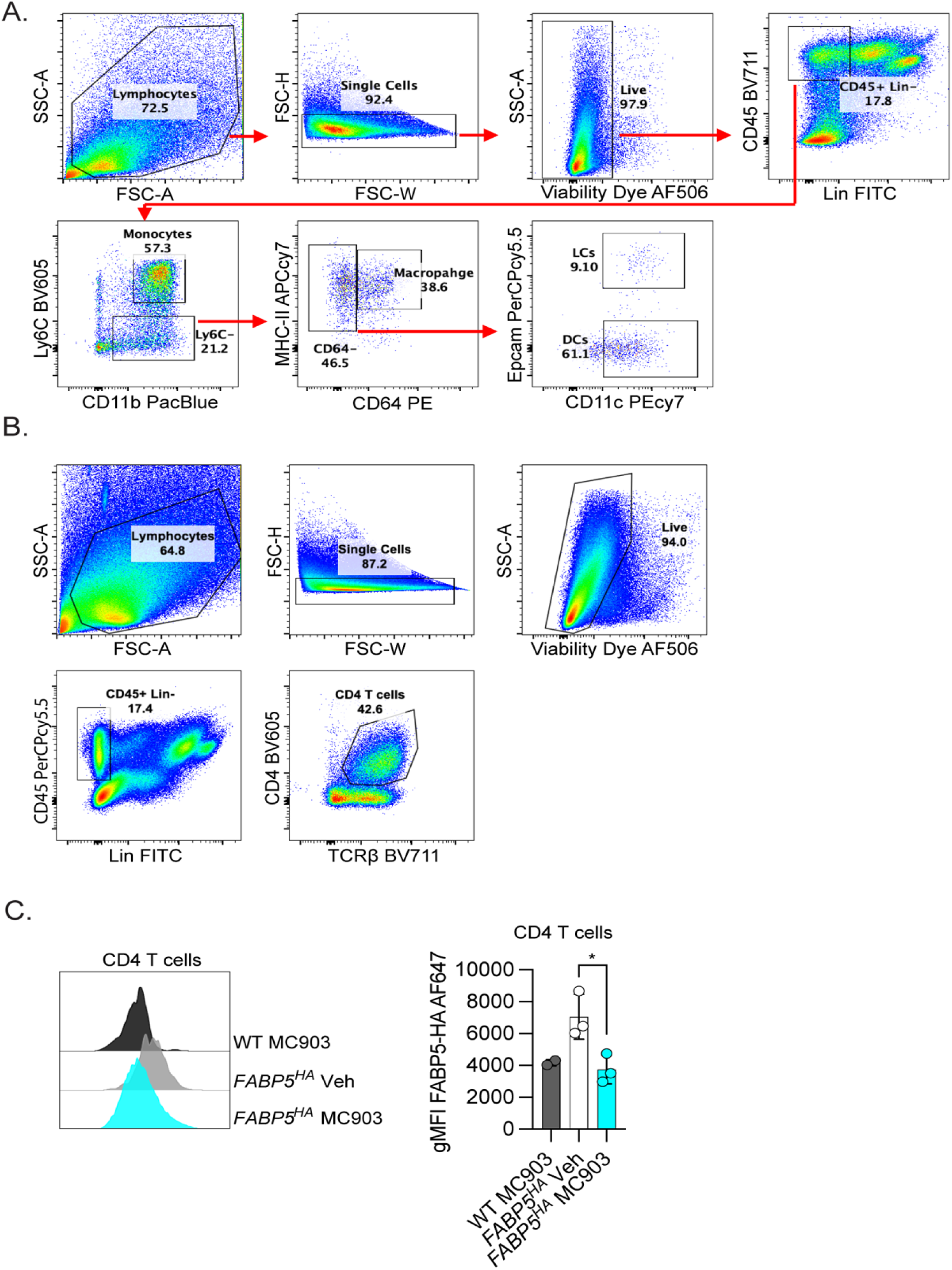
A) Flow cytometry gating strategy for identification of monocytes, macrophages, dendritic cells (DCs) and Langerhans cells (LCs) for quantification of intracellular FABP5-HA AF647 fluorescent intensity related to figure 2F. B) Flow cytometry gating strategy for identification of CD4^+^ T cells for quantification of intracellular FABP5-HA AF647 fluorescent intensity related to S3C. C) Histogram (left) comparing the fluorescent intensity of intracellular á-ÇÁ AF647 staining in different CD4^+^ T cells isolated from skin of WT and FABP5^HA^ mice treated with vehicle or MC903. Quantification of geometric mean fluorescent intensity (gMFI) (right) from these same populations (n = 12). Unless otherwise noted, all data are reported as means ± SD. * *p* < 0.05; ** *p* < 0.01, *** *p* < 0.005. Two-tailed unpaired Student’s *t*-test. Datapoints are discrete biological replicates. (C) and (D) were analyzed by one-way ANOVA with Fisher’s least significant difference test.

**Supplemental Figure 4:**
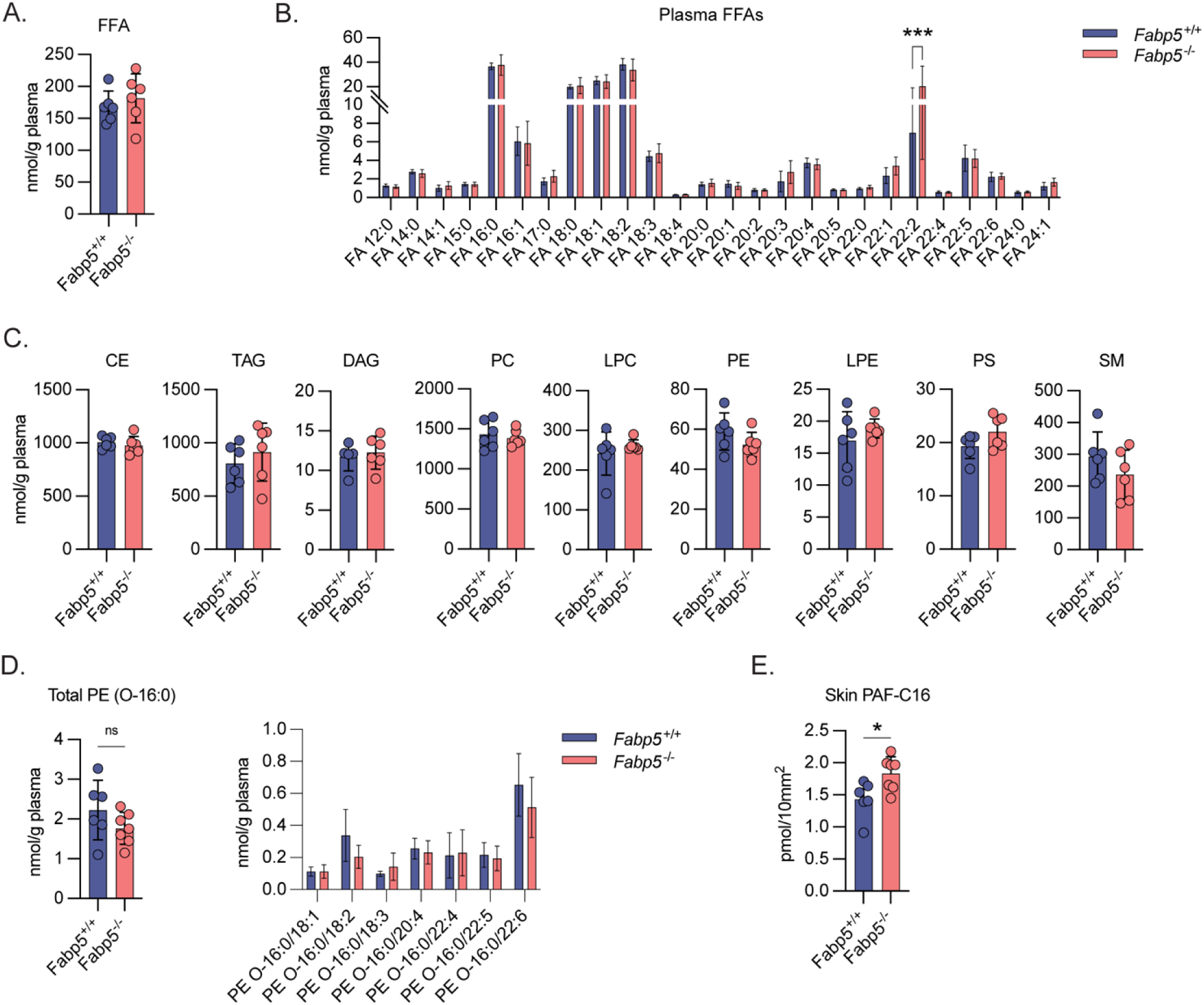
A) HPLC-MS/MS based quantification of total free fatty acids (FFA) in plasma of *Fabp5^+/+^* and *Fabp5^-/-^* mice on day 10 of treatment with MC903. B) HPLC-MS/MS based quantification of individual FFA species in plasma of *Fabp5^+/+^* and *Fabp5^-/-^* mice on day 10 of treatment with MC903. C) HPLC-MS/MS based quantification of total cholesterol ester (CE), triacylglycerol (TAG), diacylglycerol (DAG), phosphatidylcholine (PC), lysophosphatidylcholine (LPC), phosphatidylethanolamine (PE) lysophosphatidylethanolamine (LPE), phosphatidylserine (PS), and sphingomyelin (SM) species in plasma of *Fabp5^+/+^* and *Fabp5^-/-^* mice on day 10 of treatment with MC903. D) HPLC-MS/MS based quantification of total PE species with 16:0 ether-linked tails (left) and quantification of individual PE species with 16:0 ether-linked tails (right) in plasma of *Fabp5^+/+^* and *Fabp5^-/-^* mice on day 10 of treatment with MC903. E) Targeted LC-MS/MS based quantification of PAF C-16 from skin of *Fabp5^+/+^* and *Fabp5^-/-^* mice on day 10 of treatment with MC903. Unless otherwise noted, all data are reported as means ± SD. * *p* < 0.05; ** *p* < 0.01, *** *p* < 0.005. Two-tailed unpaired Student’s *t*-test. Datapoints are discrete biological replicates. (B) and (D) were analyzed by two-way ANOVA with Šidák correction for multiple comparisons.

**Supplemental Figure 5:**
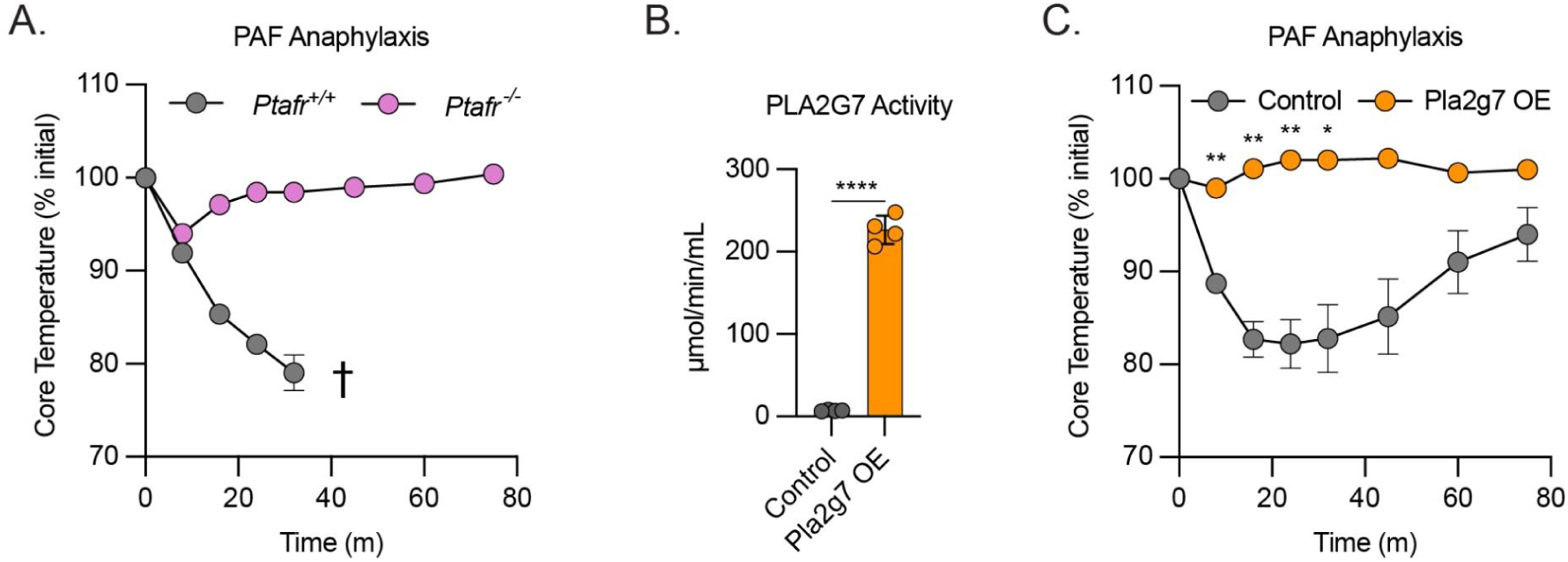
A) Core body temperatures (% of initial) following PAF induced anaphylaxis in naïve *Ptafr^+/+^* and *Ptafr^-/-^* mice. (n = 6). Dots show mean ± SEM. Stars represent Šidák corrected *p* values for 2-way ANOVA with multiple comparisons. B) PLA2G7 enzymatic activity on 2-thio PAF from serum of mice one day following hydrodynamic injection with control plasmid or plasmid encoding Pla2g7 (PLA2G7 OE). (n = 8) C) Core body temperatures (% of initial) following PAF induced anaphylaxis in mice one day following hydrodynamic injection with plasmid encoding Pla2g7 (PLA2G7 OE) or equal volume saline control. (n = 8). Dots show mean ± SEM. Stars represent Šidák corrected *p* values for 2-way ANOVA with multiple comparisons. Unless otherwise noted, all data are reported as means ± SD. * *p* < 0.05; ** *p* < 0.01, *** *p* < 0.005. Two-tailed unpaired Student’s *t*-test. Datapoints are discrete biological replicates.

